# Nonsense mutations can increase mRNA levels

**DOI:** 10.1101/2025.11.16.688500

**Authors:** Precious O. Owuamalam, Md. Nazmul Hossain, Saverio Brogna

## Abstract

Nonsense mutations are often associated with reduced mRNA levels, as premature translation termination can lead to the activation of nonsense-mediated mRNA decay (NMD). To examine how positional context influences these outcomes, we introduced premature translation termination codons (PTCs) at 15 locations within the coding region of a GFP reporter gene in *Schizosaccharomyces pombe*. PTCs in the first third of the coding region (codons 1–88) consistently led to reduced mRNA levels. In contrast, most downstream PTCs (codons 108–231) showed modest or minimal reductions, and several were associated with increased mRNA levels relative to the PTC-less control transcript. Measurement of transcript stability for one such variant indicated that the increased abundance was not attributable to decreased turnover. Deletion of *UPF1* in wild-type cells elevated the levels of transcripts that were reduced and, unexpectedly, further increased the abundance of transcripts that exceeded the control level. In spliced versions of these constructs, downstream PTCs generally reduced mRNA levels regardless of exon junction position; additionally, for some early PTCs, splicing appeared to suppress rather than enhance mRNA reduction. Overall, these observations indicate that an unexpected consequence of nonsense mutations can be increased mRNA levels. These findings may aid in the interpretation of the effects of nonsense mutations on mRNA abundance beyond the predictions of current NMD models and may also help in the design of eukaryotic gene-expression constructs.

## Introduction

Nonsense-mediated mRNA decay (NMD) is considered a quality-control mechanism of gene expression in eukaryotic cells that detects and eliminates mRNAs carrying a premature translation termination codon (PTC). The central function of NMD is therefore thought to be the elimination or the limitation of the accumulation of a broadly defined category of aberrant transcripts, which, if translated, would result in the wasteful production of truncated and potentially toxic polypeptides (1-5). NMD is also believed to regulate the expression of normal transcripts, such as those containing short upstream open reading frames (uORFs) or stop codons in an abnormal sequence context (6). This includes splice variants with poisonous exons encoding a PTC that functions to repress gene expression (7). NMD might also regulate cryptic noncoding RNAs and long noncoding RNAs (lncRNAs) because these transcripts often contain short open reading frames (8, 9). Whether NMD acts primarily as an mRNA surveillance system or as a broad gene-regulatory mechanism remains unclear, and its physiological significance is still debated across organisms (4, 10). Despite these uncertainties, NMD is of considerable medical importance. Approximately 30% of human inherited genetic diseases are caused by nonsense or frameshift mutations that introduce a PTC (11). For such allelic variants, whether the resulting transcript is subject to NMD, and the extent to which it is affected, are important determinants of disease severity and a key criterion in assessing the clinical relevance of newly identified nonsense and frameshift variants in diagnostic sequencing (12). Additionally, whether a PTC-containing transcript is affected by NMD is relevant in cancer biology, particularly as a predictor of the efficacy of cancer immunotherapy (13).

The mechanism that distinguishes premature translation termination from termination at a normal stop codon is complex and arguably remains a major unresolved question (4, 10). Many different NMD models have been proposed both within and across organisms. Two broad conceptual models have been proposed. The first is the “faux 3′UTR” model, first proposed in *Saccharomyces cerevisiae*, which posits that the key determinant of an NMD-inducing PTC is an abnormally long 3′UTR (14, 15). The other is the exon junction complex (EJC) model, first proposed in mammalian cells, which proposes that a nonsense mutation induces NMD when located sufficiently upstream of at least one splice junction (3). The key assumption of both models is that the signal that distinguishes a normal from an abnormal stop codon and induces activation of the NMD pathway is the presence of a feature downstream of the termination codon: either an abnormally long 3′UTR or the presence of an EJC (16), or the accumulation of NMD-inducing factors, such as UPF1(17). While these models can be applied as rough predictors of whether a given PTC-containing transcript will be affected by NMD, there are many observations that neither model can explain. Additionally, in many instances, nonsense mutations predicted to be subject to NMD have little or no effect on the mRNA level (4, 10, 18).

Here, we examined the effects of 15 nonsense mutations distributed throughout a GFP reporter gene on mRNA levels in *Schizosaccharomyces pombe*. GFP reporter genes have been well characterised in the context of NMD in several organisms, including *S. pombe*, a good model organism for studying NMD and testing current models (19, 20). Unlike *S. cerevisiae*, approximately 50% of its genes contain introns, and there is evidence that nonsense mutation-induced mRNA reduction often depends on splicing, but this dependence cannot be explained by an EJC-like mechanism as described for mammalian cells (19). The data unexpectedly show that the GFP coding region is largely refractory to NMD; in most cases, the presence of a nonsense mutation does not induce a reduction in mRNA level. In fact, in many instances, the mRNA level may even increase compared to the wild-type control, and for a PTC located after the protein functional domain, it also leads to an increase in GFP fluorescence. We discuss how these data might challenge and refine current assumptions about NMD and how they may also be informative when designing expression constructs.

## Results

### Most of the GFP coding region is refractory to NMD and at some positions a PTC leads to increased mRNA level

In order to extend our previous analysis of nonsense mutations on mRNA levels, we generated additional constructs at GFP codon positions 1, 40, 53, 70, 88, 108, 112, 126, 141, 161, 185, 210, and 231, totalling 15 PTCs, including PTC6 and PTC27 from our previous study (Figure 1A) (19). We integrated these constructs (where GFP is under the control of the *nmt41* promoter and *adh1* terminator) into the *leu1* chromosomal locus in a wild-type (WT) *S. pombe* strain to minimise cell-to-cell variation due to differences in plasmid copy number, as previously described (19). The constructs were extensively characterised and sequenced (Supplementary Figure 1). We observed that mutations in positions 1–88 cause a significant decrease in mRNA levels (Figure 1B). However, for PTCs located downstream from PTC108 (except for PTC126), there was no change in mRNA levels. Surprisingly, three of the mutations (PTCs 141, 185, and 231) led to increased mRNA levels. We designated Region 1 as comprising the PTCs early in the coding region, which caused reduced mRNA levels (PTC1–88), and Region 2 (PTC108– 231) as comprising mutations located further downstream, which cause no reduction in mRNA levels, or in some cases an increase in the mRNA levels (Figure 1A, C). When we compared the mean fold changes between the two regions, we observed a significant difference, further reinforcing the sharp difference in NMD sensitivity between the two regions (Figure 1C).

**Figure 1:**
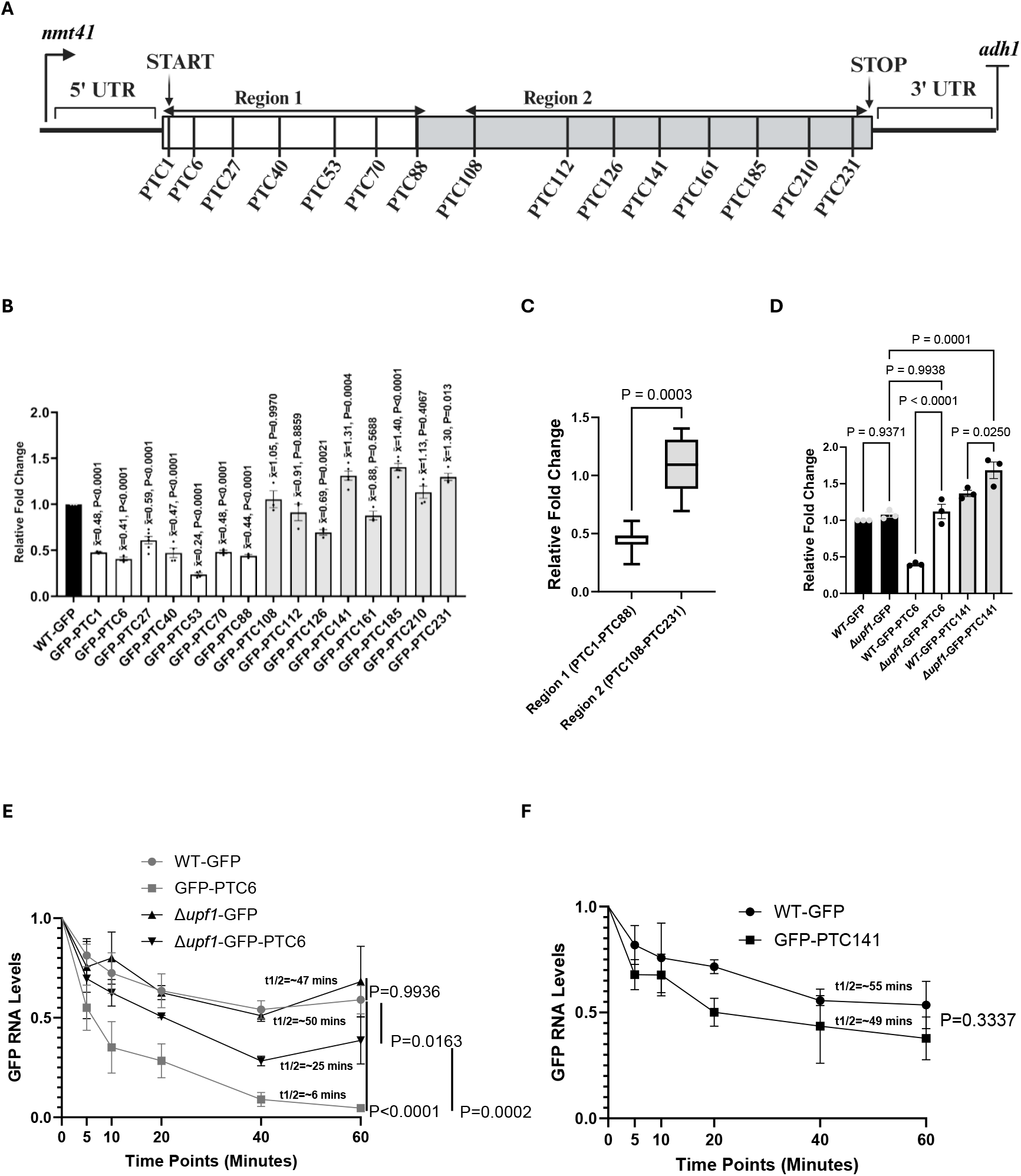
Nonsense mutations can increase as well as reduce mRNA levels. **A)** Schematic of the pDUAL-GFP construct with the nonsense mutation positions. The codon sequence is split into two regions: Region 1, where mutations reduce mRNA levels, and Region 2, where mutations either increase or do not change mRNA levels. The image was created using BioRender. **B)** qRT-PCR comparing transcript levels of the GFP-PTC-containing transcripts. All data were normalised to WT-GFP RNA levels, with *rpl32* serving as an internal control. Error bars indicate standard errors from replicate experiments (at least three biological replicates were included in all cases, as represented by the data points). A one-way ANOVA with Dunnett’s multiple comparisons was used to test for statistical significance (= mean fold change value). **C)** Cumulative comparison of the mean fold-change values of mutations in Region 1 (PTC1–88) and mutations in Region 2 (PTC108–231). Mann-Whitney rank sum test was used to test for statistical significance. **D)** qRT-PCR comparing transcript levels of WT GFP, *Δupf1-*GFP, GFP-PTC6, *Δupf1*-GFP-PTC6, GFP-PTC141, and *Δupf1*-GFP-PTC141. Transcript levels were normalised to *rpl32*, as an internal control. Error bars represent the standard error of the mean values between replicate experiments (n=3). A one-way ANOVA with Dunnett’s multiple comparisons was used to test for statistical significance. **E)** RNA stability quantifications of the WT-GFP, GFP-PTC6, *Δupf1-*GFP, and *Δupf1*-PTC6 transcripts after treatment with 1,10 phenanthroline (300 µg/mL) for the indicated time points. At least three independent biological replicates were included for each experiment. 18S rRNA was used as the internal control for qRT-PCR. For each transcript, each time point was normalised to time 0 (T0) in that transcript. Error bars represent the standard error of the mean between replicate experiments. A 2-way ANOVA with Dunnett’s multiple comparisons was used to test for statistical significance. **F)** RNA stability quantifications of the WT-GFP and GFP-PTC141 as in E. Error bars represent the standard error of the mean between replicate experiments (n=3). A t-test was used to test for statistical significance.

To examine whether the observed mRNA reduction depends on *UPF1*, we selected two representative mutations from the two regions, PTC6 (Region 1) and PTC141 (Region 2). Deletion of *UPF1* increased PTC6 mRNA levels, but unexpectedly also increased PTC141 mRNA levels, which were already significantly higher than those of PTC-less GFP (WT-GFP) (Figure 1D). We then examined the changes in RNA stability. PTC6 had lower mRNA stability (with a half-life of approximately 6 minutes) than WT-GFP (approximately 50 minutes), and deletion of *UPF1* only partially increased its stability (to approximately 25 minutes) (Figure 1E). WT-GFP and *Δupf1*-GFP were similarly stable. While the increase in GFP-PTC6 mRNA levels (Figure 1D) can partially be explained by increased RNA stability, since the deletion of *UPF1* largely restored the steady-state mRNA levels to WT levels and increased its RNA stability, that of PTC141 cannot. Although we did not test RNA stability changes between GFP-PTC141 and *Δupf1*-GFP-PTC141, we did not observe any significant changes in the stability of GFP-PTC141 (which produced more mRNA than the WT-GFP) compared to WT-GFP (Figure 1F).

### PTC231 increases GFP expression

As part of the GFP reporter characterisation, we examined the expression levels of the different reporters. As expected, the WT-GFP reporter produced detectable GFP fluorescence. In contrast, all of the PTC-containing reporters showed no detectable GFP signal, except for GFP-PTC231, which showed detectable fluorescence (Supplementary Figure 1C). Notably, its fluorescence level was approximately twice that of WT-GFP (Supplementary Figure 1D). A previous study using deletion and mutational analysis reported that the minimal domain necessary for GFP fluorescence lies between codons 7-229 (21). Therefore, codon position 231 falls outside this minimal domain required for GFP fluorescence. However, it was nonetheless surprising that this reporter produced approximately twofold more GFP fluorescence than WT-GFP given that the mRNA level was increased by only 30%.

### No evidence for a correlation between NMD resistance and translation re-initiation

Given that PTCs located in Region 2 showed little or no sensitivity to NMD, we explored potential mechanisms underlying this effect. Since translation re-initiation at a downstream ATG can prevent NMD activation (22), we considered whether such re-initiation occurs in our reporter constructs. The next methionine codon after the first ATG is located at codon position 79, nine codons away from PTC70 (Supplementary Figure 1A). Given that the level of the PTC70 transcript is increased relative to that of PTC53, it could be argued that this might be a consequence of translation re-initiation occurring at position 79. However, we considered this unlikely, since the level of PTC70 transcript is similar to that of other early PTCs. Additional methionine codons are located at positions 89, 154, 219, and 234 (Supplementary Figure 1A). When comparing transcript levels of PTCs upstream of these methionine codons with those further upstream or downstream, there is no correlation, indicating no evidence that any putative translation re-initiation mechanism significantly impacts NMD in this system.

### Splicing alters the effect of nonsense mutations on mRNA levels

Previously, we showed that splicing enhances NMD in *S. pombe* (19). To further explore how PTC mutations affect NMD depending on the intron’s position, we created intron-containing versions of the reporters and integrated the pDUAL-GFPivs-PTC-containing reporters into the *leu1* locus in WT *S. pombe* (Figure 2A). The insertion of an intron enhances NMD, because when the intron is present, not only PTCs in Region 1, but also most of those located in Region 2 show reduced mRNA levels (Figure 2C). The strongest effect is seen for the PTCs closest to the intron (PTC108ivs to PTC161ivs), while the PTCs further downstream, more distant from the intron and closer to the normal stop codon (PTC185ivs to PTC231ivs), are not reduced (Figure 2B). However, we noted a few exceptions in which splicing appears to suppress NMD rather than enhance it; PTC1 and PTC53 showed increased mRNA levels in the GFPivs reporters compared with the intronless reporters (Supplementary Figure 1E).

**Figure 2:**
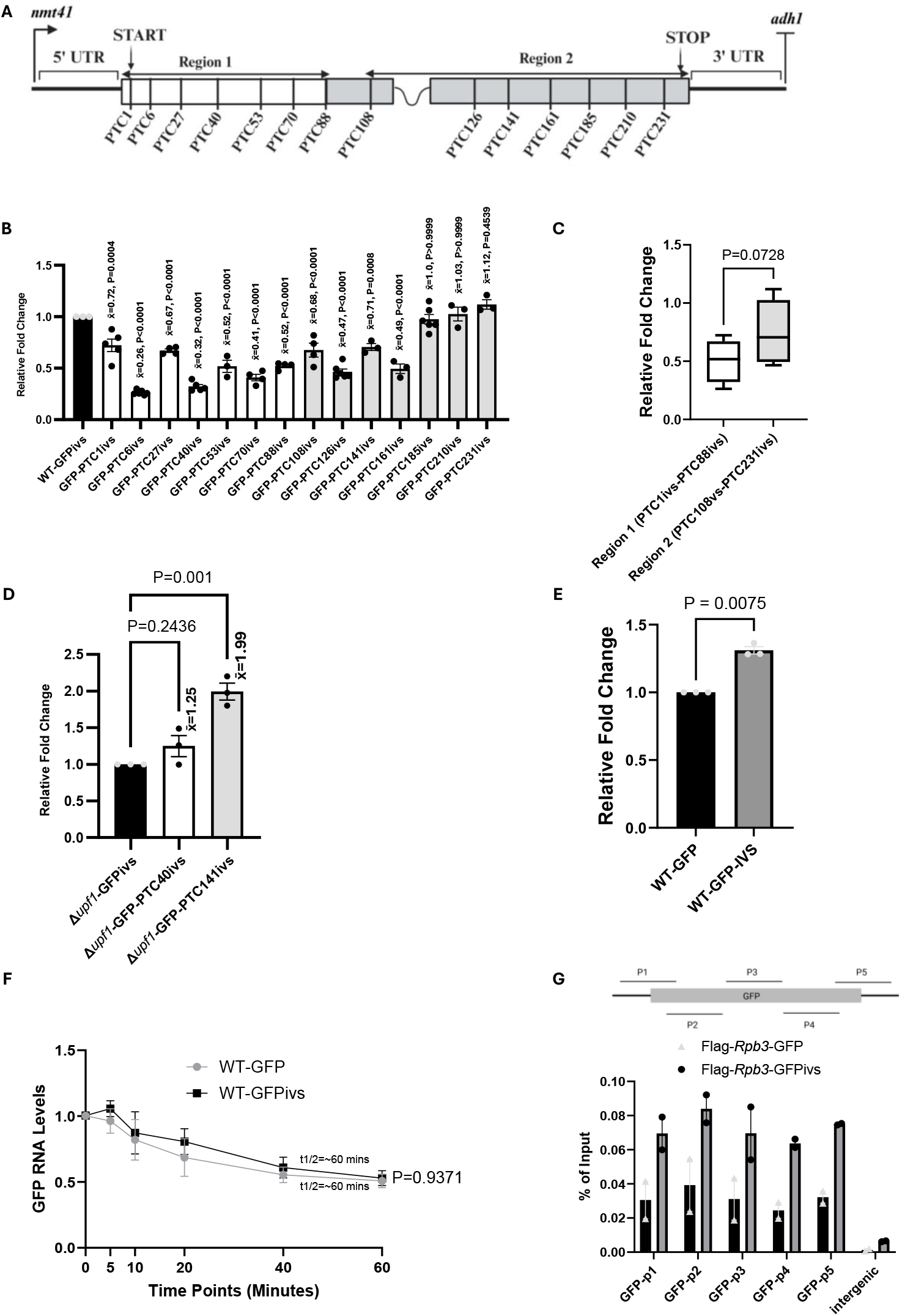
Splicing alters the effect of a nonsense mutation on mRNA levels. **A)** Schematic of the intron-containing versions of the pDUAL-GFP constructs (pDUAL-GFPivs) with the nonsense mutation positions. The codon sequence is split into two regions: Region 1, where mutations reduce mRNA levels, and Region 2, where mutations either increase or do not change mRNA levels. **B)** qRT-PCR comparing transcript levels of the GFP-PTCivs transcripts. All data were normalised to WT-GFPivs RNA levels, with *rpl32* serving as an internal control. Error bars indicate standard errors from replicate experiments (at least three biological replicates were included in all cases, as represented by the data points). A one-way ANOVA with Dunnett’s multiple comparisons was used to test for statistical significance (= mean fold change value). **C)** Cumulative comparison of the mean fold-change values of mutations in Region 1 (PTC1ivs–88ivs) and mutations in Region 2 (PTC108ivs–231ivs). Mann-Whitney rank sum test was used to test for statistical significance. **D)** qRT-PCR comparing transcript levels of *Δupf1-*GFPivs, *Δupf1-*GFP-PTC40ivs, and *Δupf1-*GFP-PTC141ivs. Transcript levels were normalised to *Δupf1*-GFPivs, with *rpl32* serving as an internal control. Error bars represent the standard error of the mean values between replicate experiments (n=3). A one-way ANOVA with Dunnett’s multiple comparisons was used to test for statistical significance (= mean fold change value). **E)** qRT-PCR comparing transcript levels of WT*-* GFP and WT*-*GFPivs. Transcript levels were normalised to WT*-*GFP, with *rpl32* serving as an internal control. Error bars represent the standard error of the mean values between replicate experiments (n=3). A t-test was used to test for statistical significance. **F)** RNA stability quantifications of the WT-GFP and WT*-*GFPivs after treatment with 1,10 phenanthroline (300 µg/mL) for the indicated time points. 18S rRNA was used as the internal control for qRT-PCR. For each transcript, each time point was normalised to time 0 (T0) in that transcript. Error bars represent the standard error of the mean between replicate experiments (n=5). A t-test was used to test for statistical significance. **G)** ChIP-qPCR ChIP signal of Pol II ChIP signal (Flag-Rpb3) on WT*-*GFP and WT*-*GFPivs (n=2). The schematic above the bar chart illustrates the approximate positions of the ChIP qPCR primers on the GFP ORF. The primer sequences are listed in Supplementary Table 2.

We examined whether the observed reductions were linked to *UPF1*. In the absence of *UPF1*, we observed a recovery of mRNA levels in Region 1 (PTC40ivs). Unexpectedly, however, we observed an about twofold increase in PTC141ivs mRNA levels above the GFPivs-PTC-less mRNA (Figure 2D), which is higher than the 1.5-fold increase seen in the intronless construct (Figure 1D). We confirmed that our qPCR primers were specific for the spliced transcripts and that we did not detect any corresponding pre-mRNA unspliced bands (Supplementary Figure 1B). Although we did not test the stability of PTC141ivs compared with WT-GFP directly, we observed an increase in mRNA levels in the GFPivs reporter compared with those of the intronless GFP reporter (Figure 2E). However, the increased mRNA levels do not correlate with an increase in mRNA stability (Figure 2F). The mechanism remains to be investigated; however, despite the reporters having the same promoter, there is more Pol II loading with the GFPivs reporter (Figure 2G), suggesting that transcripts produced from intron-containing genes are transcribed more efficiently than those without introns.

Previously, we showed that knocking out *RNPS1* or *MAGO* did not affect splicing-dependent NMD (19). We then tested whether knocking out *Y14* impacts this process (Supplementary Figure 2A). We could not test using a *Δfal1* strain because the strain is severely growth-impaired. As a control, we used the *Δmago* strain containing the WT-GFPivs, PTC40ivs, and PTC141ivs plasmids. As in the Δ*mago* strain, we did not obtain any rescue with *Δy14* (Supplementary Figures 2A and B).

Overall, the data show that splicing markedly alters the effects of nonsense mutations on mRNA levels regardless of whether the mutation is located upstream or downstream of the intron. Unexpectedly, in some instances, spliced transcripts show less nonsense-mutation-dependent mRNA reduction than the unspliced counterpart.

## Discussion

In this study, we carried out detailed mutagenesis experiments using GFP reporter constructs to examine how nonsense mutations within the GFP coding region affect mRNA levels in *S. pombe*. We found that introducing a nonsense mutation between codons 1 and 88 caused the expected reduction in mRNA levels. However, introducing a mutation between codons 108 and 231 resulted either in no reduction (apart from one instance, position 126, where a modest reduction was observed) or, surprisingly, in an increase in mRNA levels when the mutations were located closer to the natural stop codon (positions 185 to 231). It appears that the GFP coding region (239 codons in total) can be divided into two distinct regions. The first, approximately one-third of the coding sequence, is a region in which a premature translation-termination event can induce a reduction in mRNA levels. The remaining two-thirds of the coding region showed no reduction in mRNA levels. This latter portion may itself be divided into two subregions: an initial part in which premature translation termination does not alter the mRNA level, and a more distal part in which premature translation termination leads to a small increase in mRNA level. These observations are not what would be expected from the current NMD models. The constructs we are discussing lacked an intron and, therefore, were expected to follow the faux 3′UTR or similarly based NMD models. According to these, a nonsense mutation—apart from those very close to the normal stop codon—should result in a reduced mRNA level (14, 15). This reduction should be proportional to the distance from the 3′ end of the transcript and should be most apparent when the PTC is furthest from the end of the transcript. The observations confirm the previous report that lengthening the distance between the PTCs and the normal stop codon does not suppress NMD in this organism (19). These data therefore suggest that the key determinant of whether the mRNA is affected is whether premature translation termination occurs within a certain window after translation has begun. Translation is expected to be slow in the initial portion of the coding region, as is the case for other genes previously investigated (23-25). It is therefore plausible that a PTC would have an effect only during this early stage, when translation has not yet reached its optimal elongation rate, whereas once elongation has reached its optimal phase, premature termination of translation no longer affects the fate of the mRNA.

Another notable observation is that inserting a stop codon at position 1 (PTC1), most likely destroying translation initiation at the ATG start codon also resulted in a similar level of mRNA reduction. The mRNA reduction caused by PTC1 is not expected to be due to NMD, since NMD is thought to be triggered by a premature translation termination event; thus, in the absence of initiation, there would not be any termination event. It is nevertheless interesting to consider, in view of the similarity of the reduced mRNA phenotype, whether a common feature underlies both events. A shared characteristic might be that these mRNAs never underwent or did not permit the ribosome to reach its optimal rate of translational elongation. In other words, mutations that, either directly or indirectly, prevent the ribosome from reaching an optimal elongation stage render the mRNA intrinsically short-lived.

One question that arises is why a PTC located close to the normal stop codon shows an increased mRNA level in the intron-less versions of the construct. This is particularly striking given that, at least for the transcripts tested, this increase does not appear to be due to enhanced mRNA stability and may therefore reflect increased mRNA production. One possibility is that lengthening the distance between the stop codon and the polyadenylation signal can increase gene expression. This is consistent with a previous report that lengthening the distance between the normal stop codon and the polyadenylation site can increase the mRNA level of a similar GFP reporter in *S. pombe* (19). This suggests that the key determinant of gene expression that should be investigated is the distance between the stop codon and the cleavage and polyadenylation site, and possibly NMD, in light of a previous observation that nonsense mutations are linked to abnormal 3′ end cleavage and polyadenylation (26). What is also notable is that a transcript displaying elevated mRNA levels shows an even higher increase in cells depleted of *UPF1*, and the effect seems to be more apparent with the spliced construct. The data suggest that this may not be a consequence of hyper-stabilisation of the transcripts but an increase in their production. Consistent with this interpretation, previous observations indicate that many transcripts which are upregulated in *UPF1* deletion strains are also shown to have increased Pol II loading in *S. pombe* (27). The mechanism is unknown, and it remains to be determined whether it might be linked to UPF1 presence at transcription sites, as reported in both *S. pombe* and *Drosophila* (27, 28). Regardless of what the mechanism might be, this observation cautions against interpreting the upregulation of a given set of transcripts in UPF1-depleted cells as NMD targets that have been stabilised, as previously discussed (27).

Similarly to what has been reported previously in *S. pombe* and many other organisms, splicing radically increases the likelihood of a nonsense mutation leading to mRNA reduction. These data show that splicing enhances the mRNA reduction regardless of whether the intron is located upstream or downstream of the PTC, contrasting with the expectation of an EJC-like model, consistent with the earlier report in *S. pombe* (19). What the alternative mechanism might be remains unknown; some hypothetical models have been considered to explain similar observations in *S. cerevisiae* (29). Consistent with previous observations that splicing enhances gene expression (30, 31), we noticed that the presence of an intron enhances the mRNA expression levels. However, this does not seem to be due to stability, which remains unchanged in our case. Instead, it is likely driven by an unidentified mechanism. Surprisingly, in some instances, splicing can also have the opposite effect of suppressing the NMD phenotype observed with the intronless construct (for example, PTC53 vs PTC53ivs and PTC1 vs PTC1ivs) (Supplementary Figure 1E).

In summary, contrary to a prediction of the view that cells have an effective mechanism that detects PTC-containing mRNAs as aberrant transcripts and destroys them by a specialised mechanism, we found that nonsense mutations often leave the mRNA level unchanged and, in some instances, can even increase mRNA levels without altering their stability. This implies that nonsense mutations can potentially increase mRNA production, perhaps by incidentally placing the stop codon at a more optimal distance from the polyadenylation site in the reporter gene. As for the reduction in mRNA level seen with the nonsense mutations in the early portion of the coding region, this may be primarily triggered by the ribosome detaching from the mRNA before it has reached the optimal elongation stage, and it is the absence of having achieved optimal translation speed that signals for mRNA reduction. The proposed model is that NMD may primarily be a consequence of a failure to establish an optimal translation circuit on the mRNA. Conceptually, this would be similar to the previously proposed ribosome release model of NMD (4, 10), but with the emphasis that an optimally translated mRNA is not necessarily the one with the highest ribosomal load, as recently discussed (24). Notwithstanding the limitations of this study, as it is based on observations from a single reporter gene and a single organism, the conclusions should be relevant when designing eukaryotic expression constructs and when interpreting the phenotypic consequences of nonsense and frameshift mutations more broadly.

### Materials and Methods Yeast strains

The list of *S. pombe* strains used in this study is shown in Supplementary Table 1.

### Plasmid construction and integration

The plasmids used in this study were derived from the pDUAL vector (32). All plasmid manipulations were performed on the pDUAL-GFP plasmid (19). The different PTC mutations at codon positions 1, 40, 53, 88, 108, 112, 126, 161, 185, 210, and 231 were introduced into the pDUAL-GFP plasmid using site-directed mutagenesis (33, 34) with some modifications. We used the Q5 DNA polymerase enzyme (New England Biolabs), along with primer pairs that have non-overlapping sequences at their 3′ end and complementary, overlapping sequences at the 5′ end. The PTC mutations were placed in the overlapping complementary sequences. We chose a minimum of 6 bases upstream and downstream of the desired codon mutations to correspond to the primer’s complementary (overlapping) regions, with the primers terminating in a guanine or a cytosine. Our mutagenesis primers were about 50–60 bases in length, and the minimum melting temperature (T_m_) difference between the non-overlapping and overlapping regions was 3°C. The non-overlapping regions were designed to be longer and of higher T_m_ than the overlapping regions. Using these parameters, a two-stage PCR protocol was performed in a single run for each mutation. The first stage involved a standard PCR cycle: initial denaturation at 98°C for 30 seconds (1 cycle), followed by 30–40 cycles of denaturation at 98°C for 10 seconds, annealing at a temperature dependent on the non-overlapping sequence for 30 seconds, extension at 72°C for 30 seconds per kilobase of plasmid, and a final extension at 72°C for 10 minutes. The second stage included an optional denaturation at 98°C for 10 seconds, followed by 1–3 cycles of annealing at a temperature dependent on the overlapping sequence, and a final extension at 72°C for 10 minutes. All mutations were confirmed by Sanger sequencing. The amplified plasmids were then treated with DpnI to destroy the methylated template plasmid before integration. To generate the intron-containing versions of the plasmids, the pDUAL-GFP plasmids with the nonsense mutations were digested with Pml1 restriction endonuclease, and the second intron of *ubc4* was amplified from *S. pombe* genomic DNA and cloned into the cut site at codon 110 using the NEBuilder HiFi DNA assembly kit (New England Biolabs) following the manufacturer’s instructions. To generate the NMD reporter strains, the plasmids were integrated into the *leu1* locus as previously described (19, 32) in the WT, *Δupf1, Δmago, Δy14*, and the Flag-Rpb3 tagged strains. The list of primers used in this study is shown in Supplementary Table 2.

### RNA extraction, analysis, and RNA stability quantification

Total RNA was extracted using the hot acid-phenol method (35). After extraction, the total RNA was treated with DNase I (1 unit) (Thermo Scientific) according to the manufacturer’s instructions. First-strand cDNA synthesis was performed with the FastGene Scriptase II cDNA synthesis kit (Nippon Genetics) using 50 ng of total RNA, according to the manufacturer’s instructions. Quantitative real-time PCR assays were conducted in 96-well plates using the ABI Prism^TM^ SDS 7000 real-time PCR thermocycler (Applied Biosystems), following the manufacturer’s instructions. PCR assays were performed using the qPCRBIO SyGreen Blue Mix Hi-ROX (PCR Biosystems). The 2^-ΔΔCT^ method was used to calculate the relative expression levels of the target transcripts, normalised to *rpl32* mRNA or 18S rRNA. To inhibit transcription, cells were cultured to OD_600_ (∼0.7) in PMG media and treated with 300 μg/mL 1,10-phenanthroline (Sigma). At different time points (0, 5, 10, 20, 40, and 60 minutes) immediately after adding the drug, 10 mL aliquots were transferred into 50 mL Falcon tubes containing about one volume of crushed ice before RNA extraction. mRNA decay rates were determined by fitting an exponential decay model to transcript abundance over time using the formula: (ln(y) = ln(A) − kt). For each time point, mRNA levels were first normalised to the 0-time point value. The natural logarithm of the normalised values was then plotted against time, and a linear regression was performed, corresponding to the first-order decay model. The decay constant (*k*) was obtained from the slope of the regression line, and the mRNA half-life was calculated as: 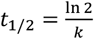.

### Chromatin immunoprecipitation (ChIP)

Freshly harvested cells from exponentially growing cultures (OD_600_ = 0.5) were fixed for 5 minutes at room temperature with 1% formaldehyde (Sigma Aldrich), followed by 10 minutes incubation with a further addition of glycine to stop the cross-linking. ChIP was then conducted as previously described (27).

### Microscopy and image quantification

Freshly harvested cells from exponentially growing cultures (OD_600_ = 0.5) were used. 1 mL of the culture was briefly centrifuged to pellet the cells. The supernatant was discarded, and the pellet was washed in 1 mL of sterile H_2_O. After a brief second centrifugation to pellet the cells, the supernatant was discarded again, and the pellet was resuspended in sterile water. A 5 μL aliquot of the cell suspension was transferred to a slide with a coverslip, prepared for viewing under the Nikon Eclipse Ti fluorescence microscope. DIC images were captured with an exposure time of 100 ms, while GFP fluorescence (FITC) was visualized with an exposure time of 5 s using immersion oil objectives. Image quantification was performed with ImageJ, and total cell fluorescence was calculated using the formula CTCF = Integrated Density – (Area of the selected cell × Mean background fluorescence).

### Statistical analysis

Statistical analysis was performed using GraphPad Prism^TM^, which was also utilized to generate the figures. A t-test was employed to compare significant differences between the control and experimental groups when the data (of not more than two groups) were normally distributed. When the data were not normally distributed, the Mann-Whitney Rank Sum test was used. For more than two data groups, ANOVA statistics were followed by Dunnett’s multiple comparisons. Statistical significance is indicated with absolute P values.

## Acknowledgement

We thank Hannah Dixon for critically reading the first draft of the manuscript. The authors acknowledge funding from the Biotechnology and Biological Sciences Research Council [BB/M022757/1] (to SB), The Darwin Trust of Edinburgh (to POO), and Bangabandhu Science and Technology Fellowship Trust (to MNH).

---

**Supplementary Figure 1:**
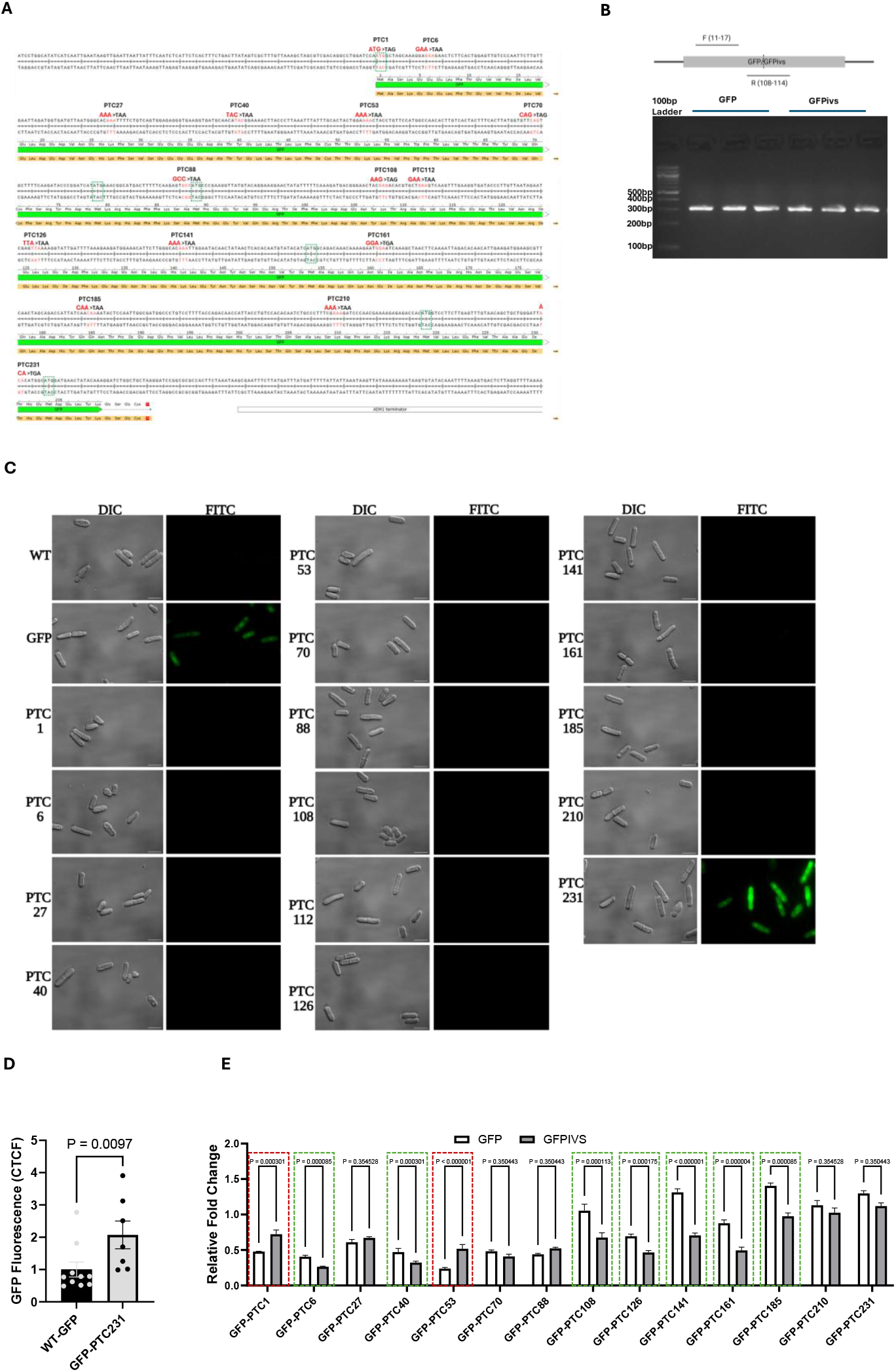
Characterisation of reporters. **A)** GFP coding sequence. Nonsense codon positions are indicated in red. The codons in each case were changed to a PTC using site-directed mutagenesis. The original codons and their corresponding PTC substitution are shown. The sequences highlighted in the green-dotted boxes show the ATG codons in GFP. The image was generated in SnapGene Viewer. **B)** Validation of the NMD qRT-PCR primers. PCR was performed under saturating conditions (40 cycles) to increase the chance of detecting any pre-mRNA or unspliced transcripts using three independent WT-GFP and WT-GFPivs cDNAs, and to verify that the qRT-PCR primers (GFP-qPCR-F and GFP-qPCR-R, sequences in Supplementary Table 2) were specific for the spliced transcript (in this case, GFP). Both primer pairs work with GFP and GFPivs, since during splicing the intron is removed, producing only spliced GFP with a cDNA similar to that of WT-GFP. The approximate positions of the qPCR primers in GFP are shown. The dotted line in the schematic above the gel signifies the splice site. A 1% agarose gel was used. **C)** Representative DIC and fluorescence (FITC) images of the WT, GFP, and GFP-PTC-containing strains. Scale bars = 10µm. **D)** Image quantification of the GFP and GFP-PTC231 cells from C. 10 WT-GFP and 7 GFP-PTC231 independent cultures were imaged, and the corresponding images were quantified using ImageJ to compare GFP fluorescence. The mean fluorescence values are shown. Error bars represent the standard error of the mean across replicates. In each image, four individual cells were quantified, totalling 40 cells for WT-GFP and 28 for GFP-PTC231. Mann-Whitney rank sum test was used to test for statistical significance. **E)** Pairwise comparison of GFP and GFPivs transcript levels. Error bars represent the standard error of the mean across replicates (n≥3). The pairwise comparisons highlighted in red represent instances where a significant increase in mRNA levels was observed in the spliced constructs compared to the intronless constructs. The comparisons highlighted in green represent instances where splicing enhances NMD. The unhighlighted comparisons indicate cases where no significant differences were observed. A t-test was used to test for statistical significance.

**Supplementary Figure 2:**
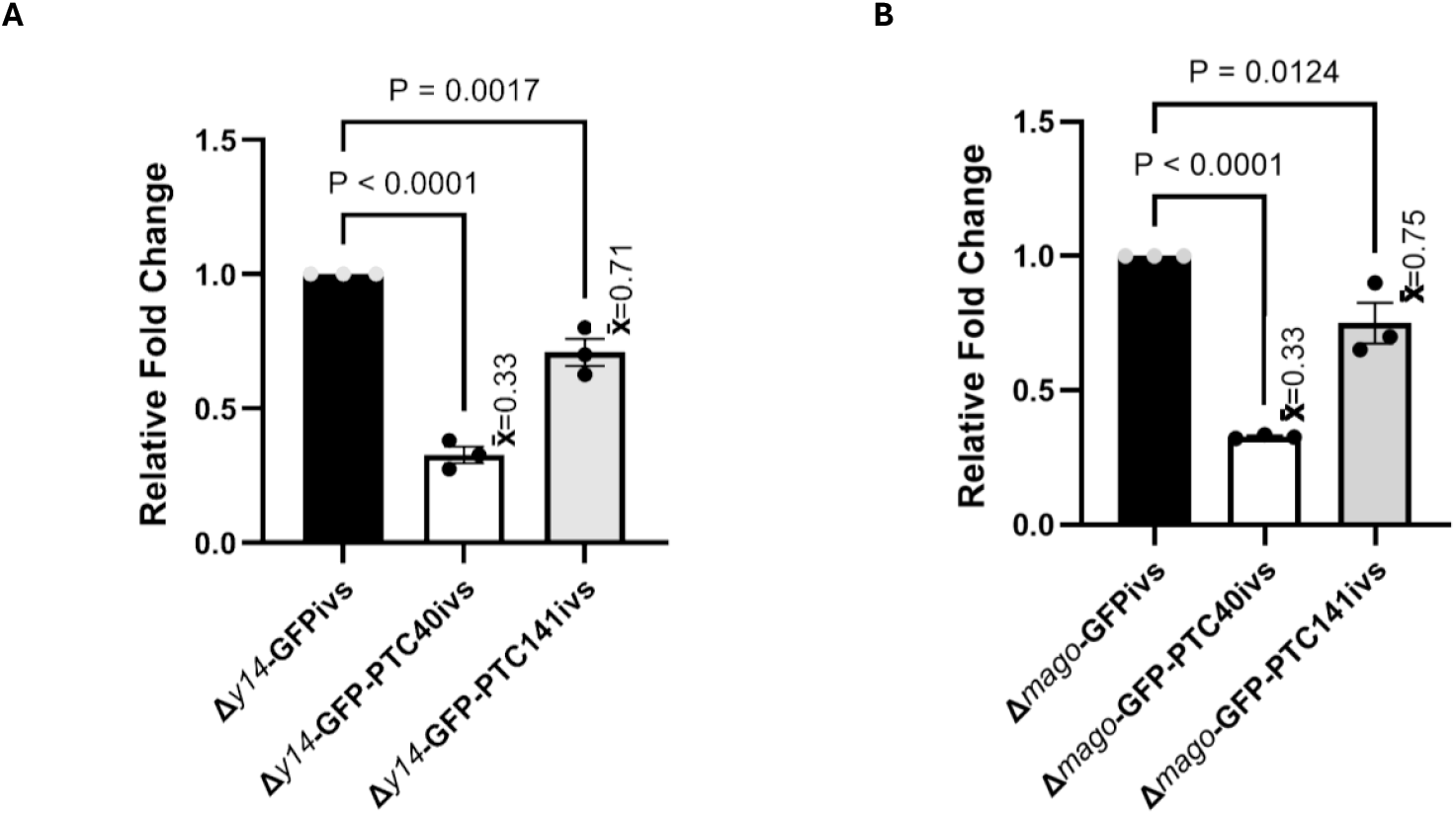
Splicing-dependent NMD does not require *Y14* and *MAGO*. **A)** qRT-PCR comparing transcript levels of *Δy14-*GFPivs, *Δy14-*GFP-PTC40ivs, and *Δy14-*GFP-PTC141ivs. Transcript levels were normalised to *Δy14*-GFPivs, with *rpl32* serving as an internal control. Error bars represent the standard error of the mean values between replicate experiments (n=3). A one-way ANOVA with Dunnett’s multiple comparisons was used to test for statistical significance (= mean fold change value). **B)** qRT-PCR comparing transcript levels of *Δmago-*GFPivs, *Δmago-*GFP-PTC40ivs, and *Δmago-* GFP-PTC141ivs. Transcript levels were normalised to *Δmago*-GFPivs, with *rpl32* serving as an internal control. Error bars represent the standard error of the mean values between replicate experiments (n=3). A one-way ANOVA with Dunnett’s multiple comparisons was used to test for statistical significance (= mean fold change value).

**Supplementary Table 1:**
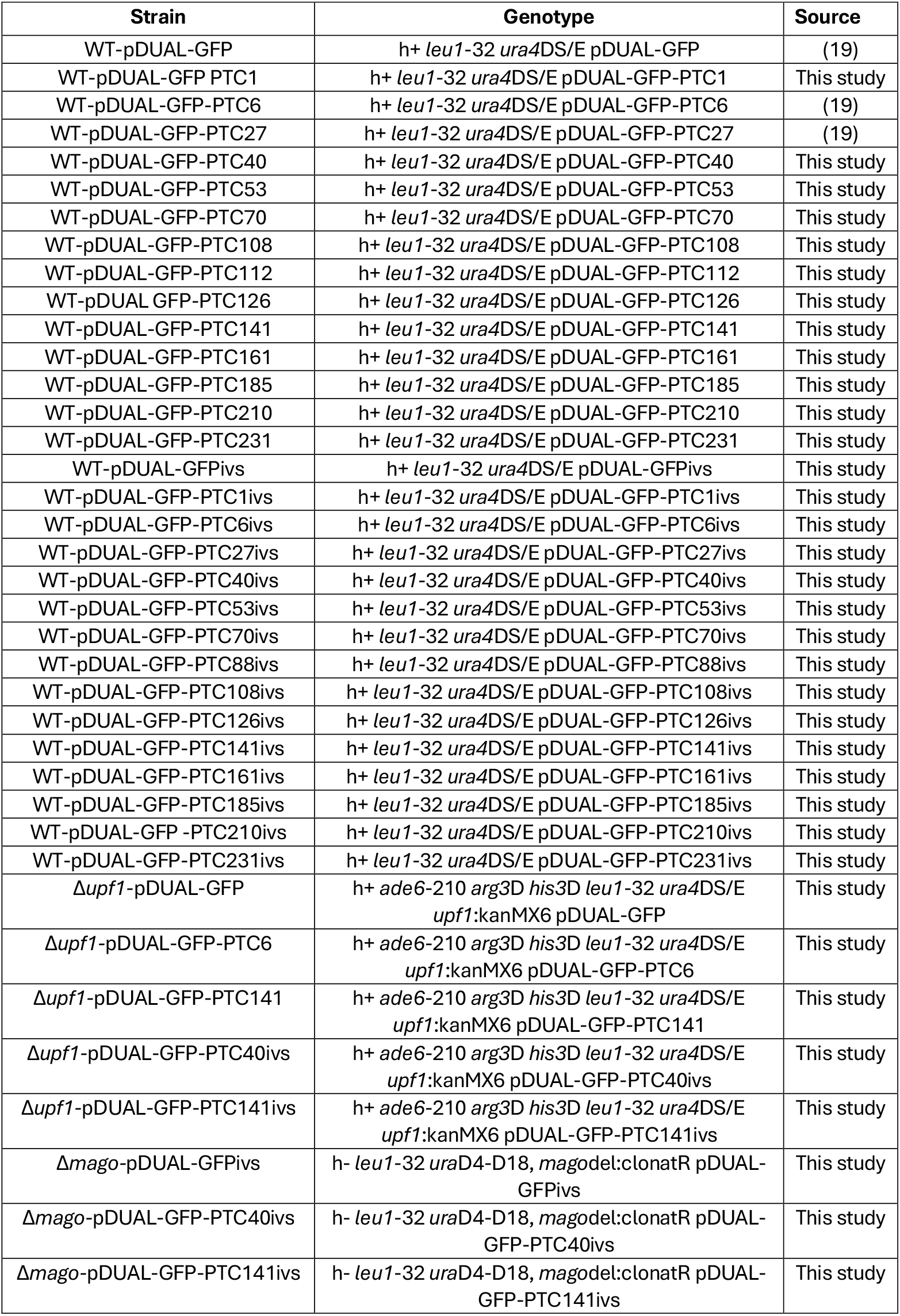
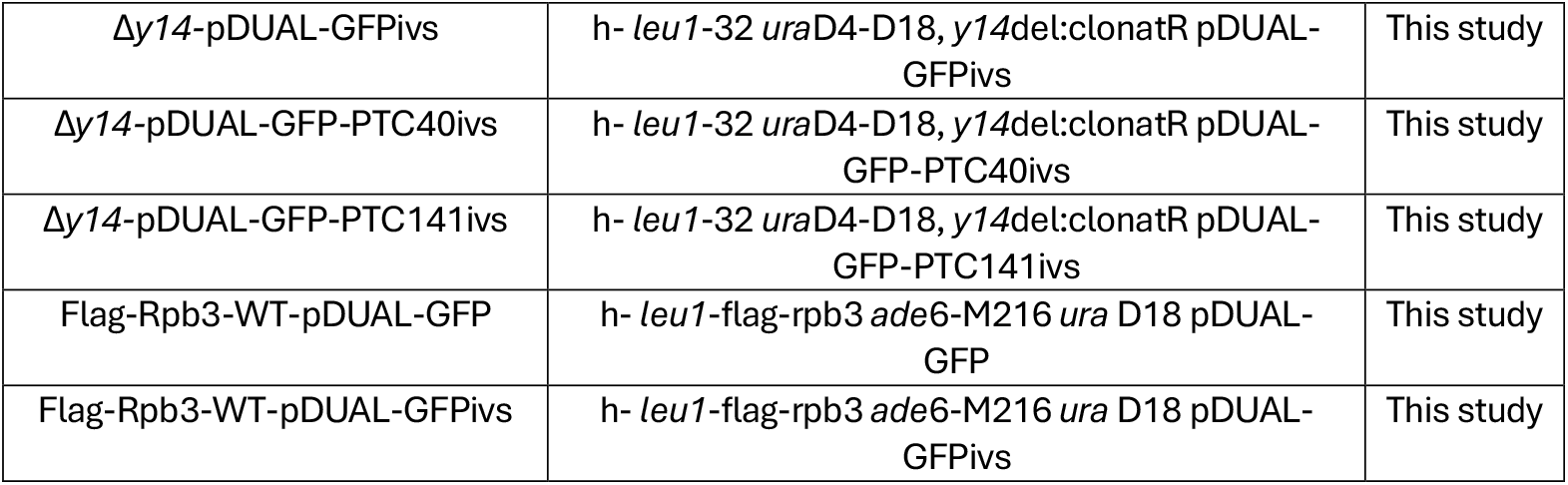
Yeast strains used in this study.

**Supplementary Table 2:**
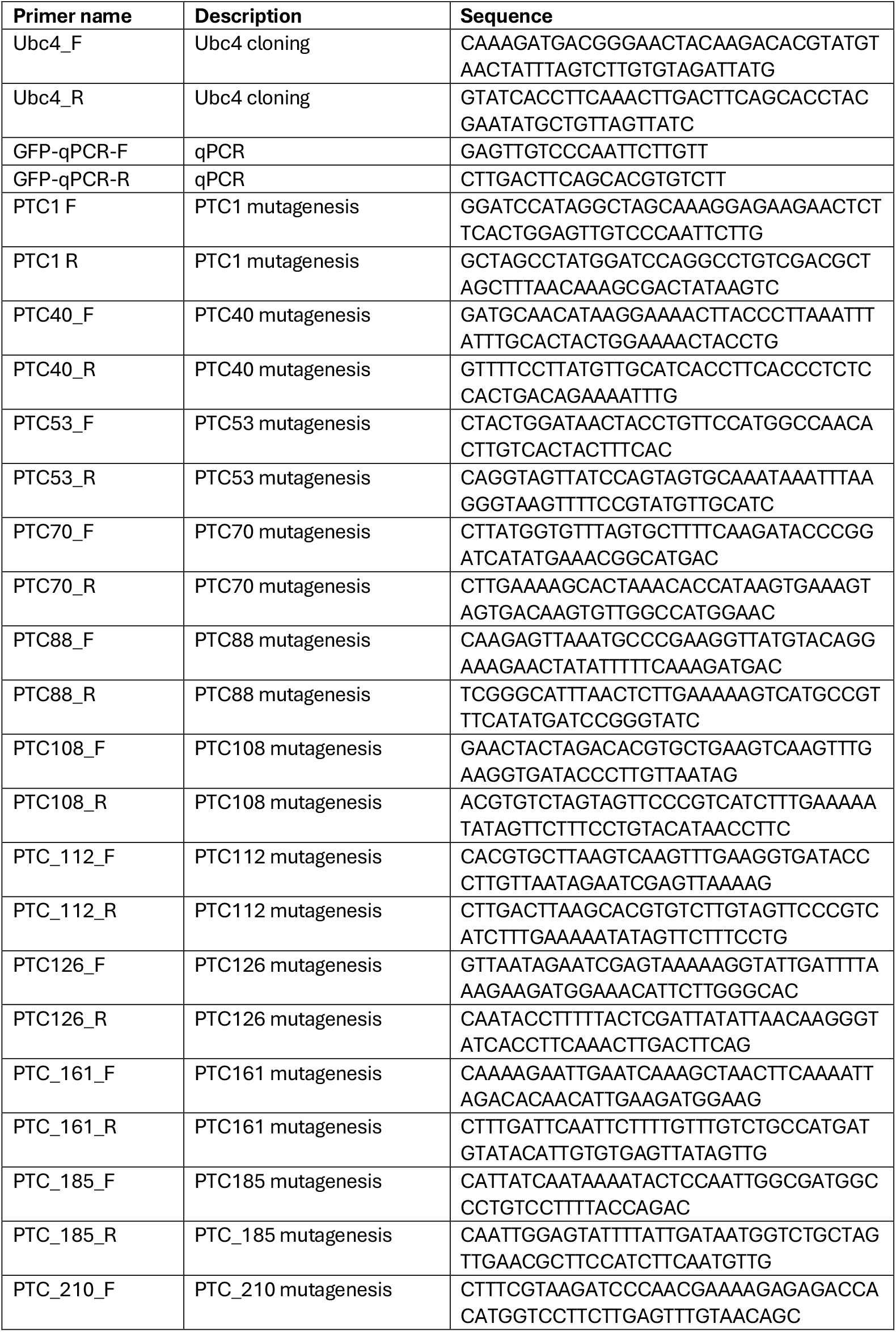
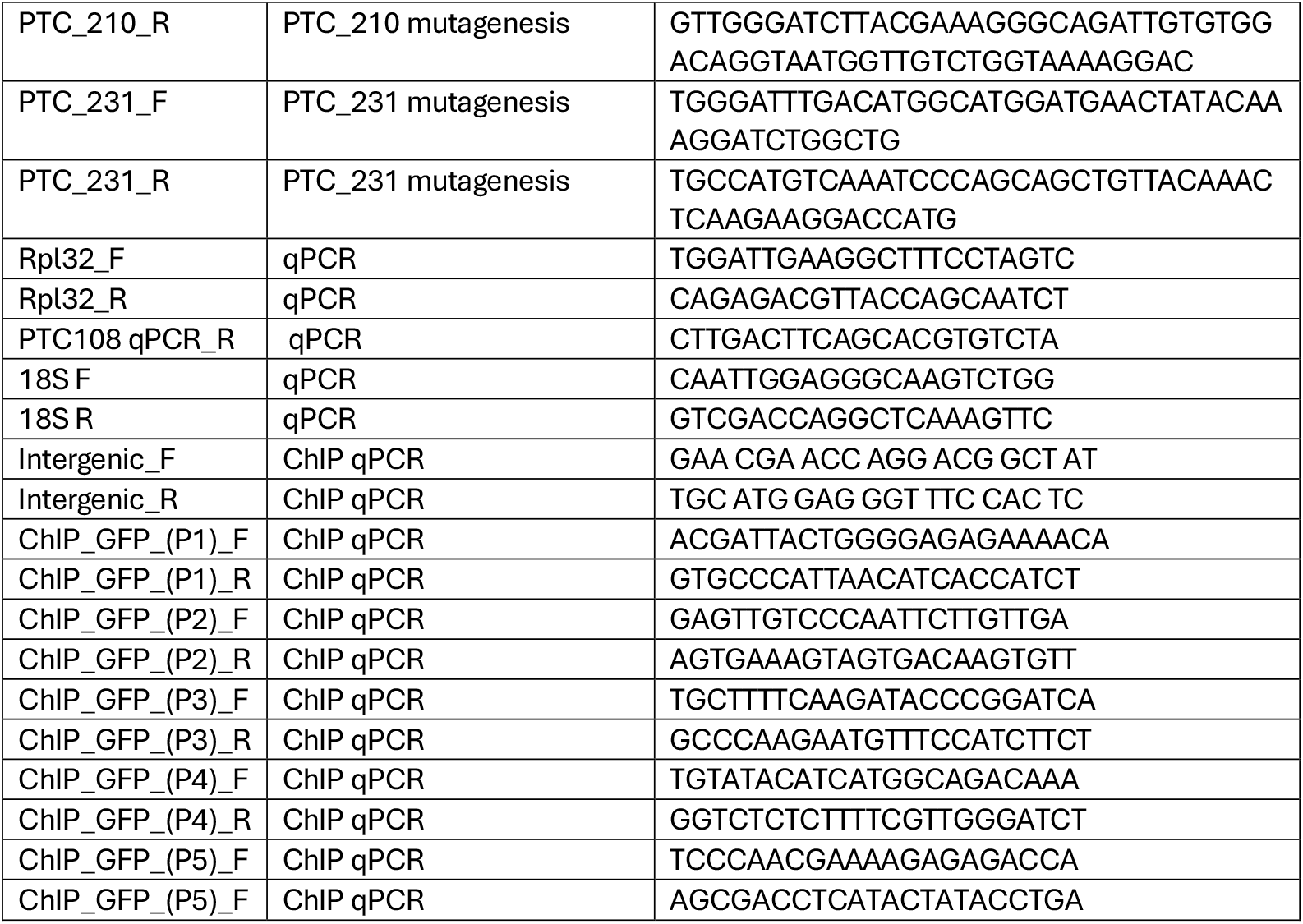
Primers used in this study.

